# Live Cell Imaging of Bioorthogonally Labelled Proteins Generated With a Single Pyrrolysine tRNA Gene

**DOI:** 10.1101/161984

**Authors:** Noa Aloush, Tomer Schvartz, Andres I. König, Sarit Cohen, Eugene Brozgol, Benjamin Tam, Dikla Nachmias, Oshrit Ben-David, Yuval Garini, Natalie Elia, Eyal Arbely

## Abstract

Genetic code expansion enables the incorporation of non-canonical amino acids (ncAAs) into expressed proteins. ncAAs are usually encoded by a stop codon that is decoded by an exogenous orthogonal aminoacyl tRNA synthetase and its cognate suppressor tRNA, such as the pyrrolysine synthetase/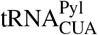 pair. In such systems, stop codon suppression is dependent on the intracellular levels of the exogenous tRNA. Therefore, multiple copies of the tRNA^Pyl^ gene (PylT) are encoded to improve ncAA incorporation. However, certain applications in mammalian cells, such as live-cell imaging applications, where labelled tRNA contributes to background fluorescence, can benefit from the use of less invasive minimal expression systems. Accordingly, we studied the effect of tRNA^Pyl^ on live-cell fluorescence imaging of bioorthogonally-labelled intracellular proteins. We found that in COS7 cells, a decrease in PylT copy numbers had no measurable effect on protein expression levels. Importantly, reducing PylT copy numbers improved the quality of live-cells images by enhancing the signal-to-noise ratio and reducing an immobile tRNA^Pyl^ population. This enabled us to improve live cell imaging of bioorthogonally labelled intracellular proteins, and to simultaneously label two different proteins in a cell. Our results indicate that the number of introduced PylT genes can be minimized according to the transfected cell line, incorporated ncAA, and application.

## Introduction

Genetic code expansion technology enables the site-specific incorporation of dozens of non-canonical amino acids (ncAAs) into proteins expressed in live organisms^1–10^. Current methodologies generally involve the use of an aminoacyl tRNA synthetase (aaRS)/tRNA pair that can facilitate the co-translational incorporation of a supplemented ncAA into a protein of interest in response to a specific codon, typically, the amber stop codon, UAG^11–13^. The aaRS/tRNA pair is referred to as an ‘orthogonal pair’ given how it should decode the specific codon without being affected by or interfering with the host cell’s translational machinery (refer to the General Introduction section in the Supplementary Information file, for a more detailed explanation). Early studies of ncAA incorporation into proteins expressed in cultured mammalian cells utilized orthogonal aaRS/tRNA pairs of bacterial origin, such as *Escherichia coli* (*E. coli*) tyrosyl tRNA synthetase and either *Bacillus stearothermophilus* or *E. coli* tRNA^Tyr^^2, 3, 14, 15^. Nowadays, the archeal pyrrolysyl tRNA synthetase (Pyl-RS) and its cognate amber suppressor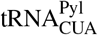^16,17^ are one of the frequently used orthogonal pairs for introducing ncAAs into proteins in cultured mammalian cells^4,18.^

Significant efforts were devoted to the development of methods for expanding the genetic code of cultured mammalian cells^2–4,19–29^. However, the experimental systems employed were based on different orthogonal aaRS/tRNA pairs, promoters, and terminators. In addition, the numbers of encoded tRNA genes and plasmids, as well as DNA delivery methods, were not identical, making it difficult to compare the results of such studies (Supplementary Table S1). That said, these studies significantly improved ncAA incorporation and protein expression levels in mammalian cells. In particular, it was found that the intracellular level of suppressor tRNA is a limiting factor in stop codon suppression efficiency and as such, in protein expression levels. Moreover, it was demonstrated that high levels of prokaryotic tRNA transcription and processing can be achieved using constitutive RNA polymerase III (Pol III) promoters, such as U6 or H1 promoters that have no downstream transcriptional elements^3,4, 20, 22, 24^. Consequently, in the majority of current systems used for genetic code expansion in cultured mammalian cells, multiple copies of ‘tRNA cassettes’ comprising the U6 and/or H1 promoter followed by a suppressor tRNA are encoded in tandem and/or on different plasmids^26–28,30^. In addition, intracellular levels of foreign tRNA, such as tRNA^Pyl^, can be elevated by stabilizing the tRNA, for example, by introducing the U25C and other mutations^24,29,31^. These studies suggest that it is crucial that the host system will be able to process the orthogonal tRNA and maintain high intracellular levels of functional tRNAs.

Proper balance between a given tRNA and its cognate aaRS is important for maintaining accurate and efficient aminoacylation, as well as for high stop codon suppression efficiency^22,32^. However, it is difficult to control intracellular levels of an aaRS and its cognate tRNA that are exogenously expressed (or transcribed) in transiently transfected cultured mammalian cells. Using a viral transfection system, it was suggested that efficient amber suppression can be achieved using a weak promoter for aaRS expression and multiple copies of the cognate suppressor tRNA gene (up to 20 copies)^30^. There are also examples of cell lines stably expressing the required genetic components created using the PiggyBac transposon system and two plasmids, each carrying 4 copies of the PylT genes^26^. While these methods offer several advantages, genetic code expansion in transiently transfected cells, where it is more difficult to fine-tune intracellular levels of aaRSs and their cognate tRNAs, is still a frequently used experimental approach.

One of the exciting applications of genetic code expansion technology is the site-specific incorporation of ncAAs carrying a functional group for subsequent live-cell chemoselective-labelling with fluorescent organic dyes^33–42^. Here, the fluorophore is covalently attached to the expressed protein using bioorthogonal chemical reactions, namely reactions in which the two reactants are capable of reacting with each other under physiological conditions, yet display only low reactivity towards other chemical groups found in living cells. Bioorthogonal labelling offers a superior alternative to the fluorescent proteins commonly used in fluorescence imaging of live cells^34,43^. Such labelling is not limited to the N- or C-terminus of the modified protein, while the organic fluorophores are much smaller than fluorescent proteins (*~*0.5 *vs. ~*27 kDa) and, therefore, may have less effect on the structure and cellular function of the labelled protein, especially small or polymeric proteins. In addition, organic fluorophores often have excellent photophysical properties which may allow for prolonged live-cell fluorescence imaging with reduced phototoxicity. In recent years, bioorthogonal labelling of proteins *via* the inverse-electron-demand Diels-Alder reaction has been demonstrated with different ncAAs carrying strained systems (i.e. alkene or alkyne) and tetrazine-conjugated organic fluorophores^36–38,40, 44^.

Nontheless, live-cell imaging of bioorthogonally labelled intracellular proteins using *cell-permeable* fluorophores remains a challenge. Earlier attempts suffered from high background fluorescence observed upon addition of cell-permeable fluorophores, such as tetrazine-conjugated fluorophores^40,42^. One notable source for this background fluorescence, other than the possible incorporation of the ncAA in response to endogenous amber codons, is the reaction between conjugated fluorophores and aminoacylated tRNAs that leads to the nuclear accumulation of fluorescently-labelled tRNAs, i.e., tRNAs charged with ncAAs that then formed a covalent bond with the tetrazine-conjugated fluorophore^40,42^. While small cell-permeable fluorophores can be readily washed out of cells, labelled tRNAs are not cell-permeable and may thus persist in live cells for hours.

Several studies in transiently transfected cells found positive correlation between the number of encoded suppressor tRNA genes, tRNA levels, and amber suppression efficiency^4,22, 27, 45^. At the same time, live-cell fluorescence imaging of bioorthogonally labelled intracellular proteins can benefit from the reduced background fluorescence associated with aminoacylated tRNAs labelled with organic dyes, as well as the minimal effects of the expression system on cell physiology. Therefore, we decided to study the effect of the number of encoded PylT genes (and cellular tRNA^Pyl^ levels) on the signal-to-noise ratio and labelling quality in live-cell fluorescence imaging of intracellular proteins. Using a single-plasmid-based expression system, we varied the number of encoded PylT genes and measured amber suppression efficiency, signal-to-noise ratio, and labelling specificity, as well as tRNA^Pyl^ mobility. We found that in transiently transfected COS7 cells, a single copy of PylT gene was not only sufficient for live cell fluorescence imaging, but also increased the number of cells presenting protein-specific labelling. Importantly, the use of minimal number of PylT genes resulted in significant decrease in background fluorescence in the nucleus and cytosol. Our data suggest that the number of encoded PylT genes should be adjusted for genetic code expansion in cultured mammalian cells, based on the specific application.

## Results

### Incorporation of BCN-Lys into proteins expressed in COS7 cells is not affected by the number of encoded PylT genes

We have recently demonstrated the high efficiency of a single plasmid-based system for the incorporation of ncAAs into proteins expressed by transiently transfected cultured mammalian cells^28^. In contrast to transient co-transfection with two or more plasmids, where different populations of cells may take up different combinations of plasmid DNA^46,47^, encoding all of the genetic components onto a single plasmid allows for better control over the ratio between the different genes introduced into transfected cells. Exploiting this expression system, we created a set of plasmids for expressing *EGFP^150TAG^-HA* controlled by the human elongation factor 1 α-subunit promoter (EF1α, plasmids **a–e**, Fig. 1A). The plasmids included 0 to 4 copies of the U25C mutant of the *Methanosarcina mazei* (*M. mazei*) PylT gene controlled by the constitutive U6 promoter^31^. To directly compare the expression of proteins carrying different ncAAs, plasmids **a–e** were cloned with wild type Pyl-RS so as to incorporate Nε-[(*tert*-butoxy)carbonyl]-L-lysine (Boc-Lys, **1**, Fig. 1B), as well as an evolved aaRS (BCN-RS)^36^ for the incorporation of bicyclo[6.1.0]nonyne-L-lysine (BCN-Lys, **2**, Fig. 1B). Expression of the aaRS gene was controlled by the human cytomegalovirus (CMV) immediate-early promoter.

**Figure 1.**
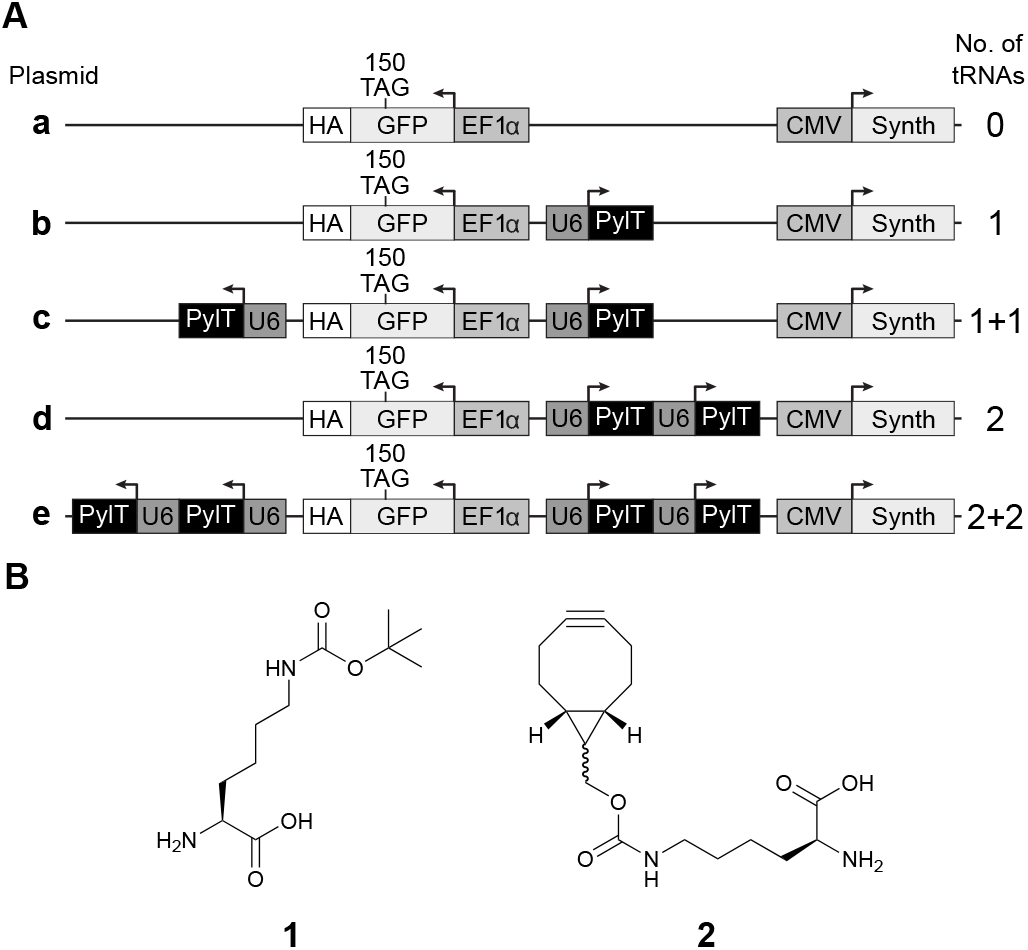
(**A**) Schematic representation of plasmids carrying different number of encoded PylT genes and *EGFP^150TAG^-HA*. Plasmid a, with no encoded tRNA served as control, while plasmid **b** carries a single copy of PylT. Plasmids **c** and **d** carry two encoded copies of PylT, either cloned separately (plasmid **c**, marked ‘1+1’) or in tandem (plasmid **d**, marked ‘2’). Plasmid **e** carries 4 copies of PylT in two cassettes, each containing two PylT genes cloned in tandem (marked ‘2+2’). One set of plasmids was cloned with wild-type Pyl-RS and a second set was cloned with evolved BCN-RS (a total of 10 plasmids). (**B**) Chemical structures of the amino acids used in this study: Boc-Lys (**1**) and BCN-Lys (**2**).

With the above plasmids, we quantified the incorporation of ncAAs **1** and **2** into 150TAG-mutant of EGFP-HA expressed in COS7 cells (Fig. 2A and Supplementary Fig. S1). The expression level of 150BocK-EGFP in cells transfected with 2*×*PylT plasmid **d** or 2+2*×*PylT plasmid **e** was higher by approximately 1.6 or 1.3, respectively, as compared to cells transfected with 1*×*PylT plasmid **b**. Although this difference is substantial, it is much lower than that measured in HEK293 cells (over 10-fold) when multiple copies of PylT were encoded in tandem^27^. Importantly, we also found that reducing the number of encoded PylT genes from 4 to 1 had a statistically insignificant effect on the expression level of 150BCNK-EGFP in COS7 cells (Fig. 2A, and Supplementary Fig. S1).

**Figure 2.**
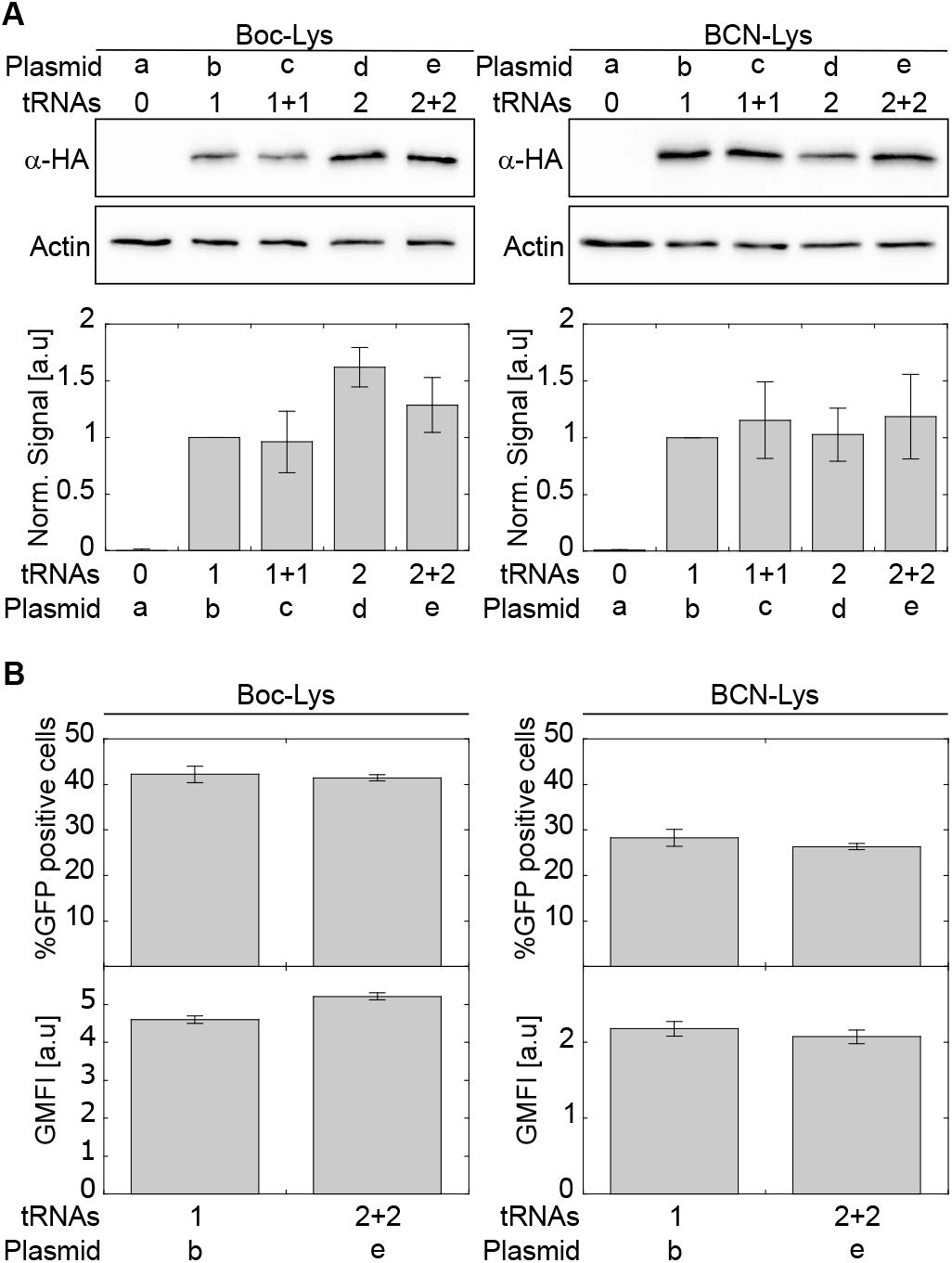
(**A**) COS7 cells were transfected with plasmids **a–e** carrying either Pyl-RS, for the incorporation of **1**, or evolved BCN-RS, for the incorporation of **2**. Expression levels of *EGFP^150TAG^-HA* were quantified by Western blot using antibodies to the C-terminal HA-tag, and normalized to actin. Expression levels as a function of indicated tRNA copy number are displayed, relative to those directed by a plasmid carrying 1*×*tRNA *±* SSD (n=5). See also Fig. S1. (**B**) COS7 cells were transiently transfected with plasmids carrying either Pyl-RS or BCN-RS and 1 or 2+2 copies of tRNA. Transfected cells were incubated with indicated ncAA for 48 h and levels of EGFP expression were analysed by FACS. The percentage of GFP-positive cells is presented as the mean *±* SSD (n=3). Protein expression levels were calculated from GFP fluorescence intensity (GFP-area) and displayed as geometric mean fluorescence intensity (GMFI*±*SSD, n=3). See also Supplementary Fig. S2 and S3.

To understand the effect of PylT copy number on live-cell imaging using BCN-Lys, we focused on plasmids carrying the minimal or maximal number of PylT genes (1*×*PylT plasmid **b** and 2+2*×*PylT plasmid **e**, respectively) and quantified resulting expression levels of the 150TAG-mutant of EGFP-HA by flow cytometry (Fig. 2B, and Supplementary Fig. S2 and S3). In excellent agreement with Western blot analyses, single-cell analysis of EGFP expression showed that reducing the number of encoded PylT genes from 4 to 1 had no effect on the percentage of viable COS7 cells expressing full length EGFP-HA, nor on protein expression levels. The data in Fig. 2B were collected 48 h post-transfection. Similar results were measured 24 h post-transfection (Supplementary Fig. S4). These data may be explained by plasmid instability, especially 2+2*×*PylT plasmid **e**, which harbors four copies of the same DNA sequence. Therefore, to check plasmid stability, we recovered the plasmids 48 h post-transfection and addressed their sizes by agarose gel electrophoresis (Supplementary Fig. S5). We found no truncations of the recovered plasmids, indicating that plasmid instability did not account for the miscorrelation between PylT copy numbers and tRNA^Pyl^ levels. Taken together, the results reveal that encoding multiple copies of PylT genes does not improve 150BCNK-EGFP expression levels in transiently transfected COS7 cells.

### Reducing the number of encoded PylT genes increases the number of specifically labelled live cells

In light of the above conclusion, we measured the effect of the number of encoded PylT genes on ncAA incorporation, in the context of bioorthogonal labelling and live-cell imaging using a cell-permeable dye. We designed four plasmids for the expression and visualization of fluorescently labelled intracellular proteins in live cultured mammalian cells as a function of encoded PylT genes (plasmids **f–i**, Fig. 3A). As a model for membrane protein labelling, we cloned *EGFP^150TAG^-CAAX* for the expression of EGFP-CAAX anchored to the inner side of plasma membrane *via* prenylation of the C-terminal CAAX motif. In addition, we cloned *HA-*α*-tubulin^45TAG^* as a model for cytoskeleton protein labelling, with the aim of labelling cellular microtubules^48^. As a fluorophore for bioorthogonal labelling we chose the cell-permeable tetrazine-conjugated silicon rhodamine (SiR-Tet) fluorophore (Supplementary Fig. S6).

**Figure 3.**
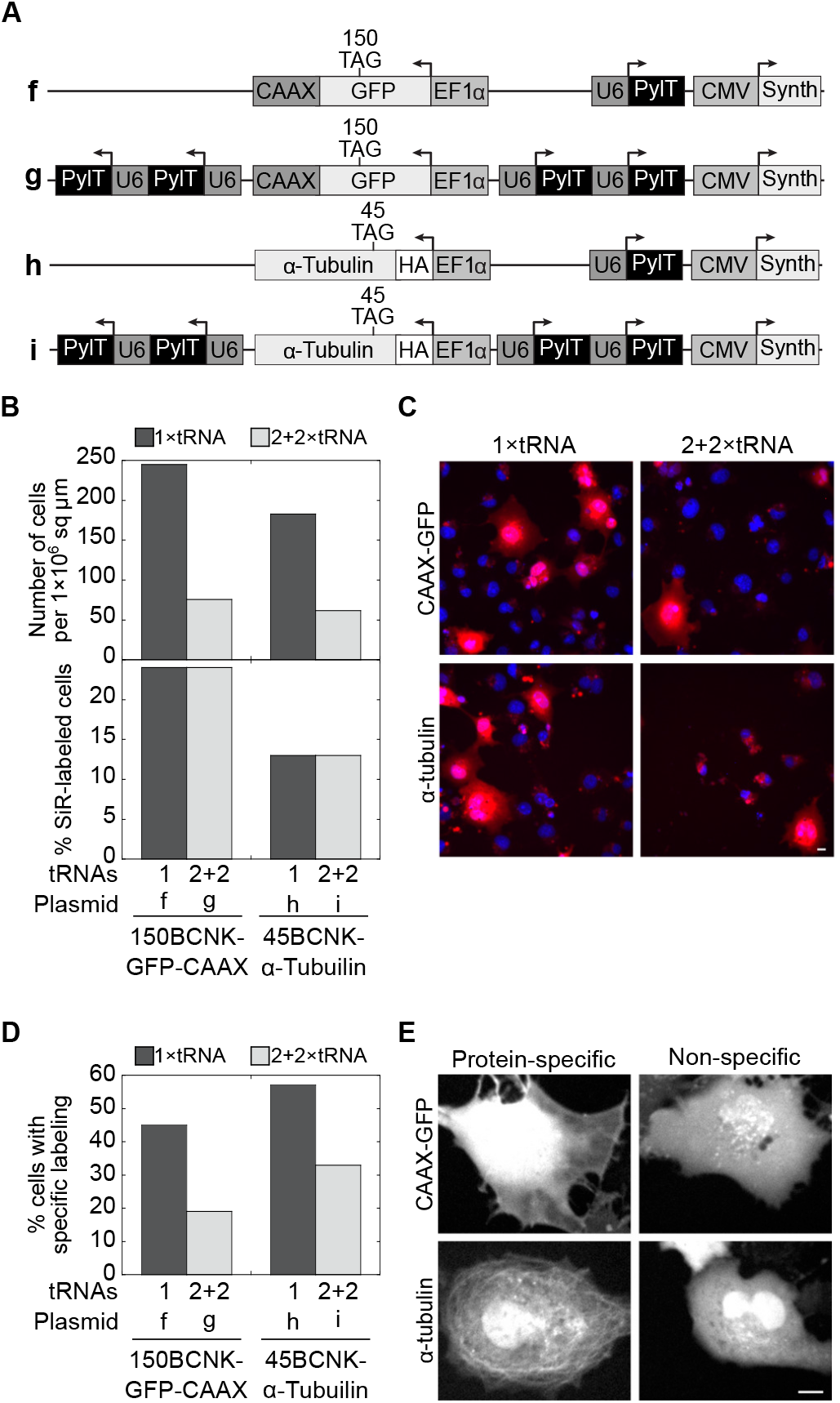
(**A**) Schematic representation of the 1*×*PylT or 2+2*×*PylT plasmids used for the expression of EGFP-CAAX (plasmids **f, g**) or α-tubulin (plasmids **h, i**), with site-specifically incorporated ncAA **2**. (**B**) Statistics of live-cell imaging of COS7 cells transfected with plasmids **f–i**, stained with Hoechst and labelled with SiR-Tet. Cells were visualized at *×*20 magnification and the total number of cells found within a 1*×*10^6^ sq µm area (top) and the percentage of SiR-labelled cells (bottom) are displayed for the indicated transfected plasmid. (**C**) Representative images of cells visualized at *×*20 magnification and quantified as described in panel B. (**D**) SiR-labelled cells (n*>*30) were visualized at *×*40 magnification and the cellular location of SiR-labelling was monitored. SiR-positive cells expressing EGFP-CAAX or α-tubulin with clear membrane labelling or labelled microtubules, respectively, were marked as cells with protein-specific labelling. The percentage of cells displaying protein-specific labelling are displayed according to the transfected plasmid. (**E**) Representative images of cells visualized at *×*40 magnification and quantified as described in panel D.

COS7 cells were transfected with 1*×*PylT plasmid **f** or **h**, or with 2+2*×*PylT plasmid **g** or **i**, incubated with ncAA **2**, and labelled with Hoechst and SiR-Tet before being visualized by live confocal microscopy. The cells that were Hoechst-positive and those that were Hoechst- and SiR-positive were counted at low magnification (*×*20, Fig. 3B top graph, and C). Within an area of 1*×*10^6^ sq µm, we found a *~*3-fold increase in the total number of Hoechst-positive cells upon reducing the number of encoded PylT genes from 4 to 1 (Fig. 3B top graph). Within the population of Hoechst-positive cells, the percentage of SiR-positive cells (i.e., cells that are both Hoechst- and SiR-positive) was independent of the number of encoded PylT genes (Fig. 3B bottom graph). This result is in agreement with flow cytometry data (Fig. 2B) showing that the percentage of GFP-positive cells was independent of the number of encoded PylT genes. However, since there were *~*3-fold more Hoechst-positive cells following transfection with the 1*×*PylT plasmid, the absolute number of SiR-positive cells was higher in samples transfected with this plasmid. In addition, using higher magnification (*×*40), we determined the localization and distribution of SiR fluorophores in single Hoechst- and SiR-positive cells (Fig. 3D and E). Within a given population of SiR-labelled cells expressing *EGFP^150TAG^-CAAX*, labelling was membrane-specific in 45% of the cells transfected with 1*×*PylT plasmid **f**, as compared to only 19% of the cells transfected with 2+2*×*PylT plasmid g. A similar trend was noted in cells expressing *HA-*α*-tubulin^45TAG^* (57% *vs*. 33%, Fig. 3D). Hence, the use of a lower number of encoded PylT genes was accompanied by an increase in the total number of live cells with protein-specific SiR-labelling.

### The use of a minimal number of encoded pylT genes improves the performance of live cell imaging

We assumed that the above described improvement in specific labelling (Fig. 3D and E) was the result of an increase in signal-to-noise ratio. To test our assumption, we performed live-cell fluorescence imaging of bioorthogonally labelled proteins using different PylT copy numbers (Fig. 4). To evaluate the signal-to-noise ratio, we measured EGFP-CAAX and SiR fluorescence intensities across the plasma membrane (yellow line, Fig. 4A, top row). When 150BCNK-EGFP-CAAX was expressed from 2+2*×*PylT plasmid **g**, a sharp peak in intensity level was measured in the GFP channel, representing membrane-anchored 150BCNK-EGFP-CAAX. At the same time, practically no notable membrane-specific increase in intensity level was obtained in the SiR channel (Fig. 4B). Indeed, SiR fluorescence intensity levels measured in the cytosol were similar to fluorescence intensity levels measured in the membrane. In contrast, when 150BCNK-EGFP-CAAX was expressed from 1*×*PylT plasmid **f**, a sharp peak in intensity levels in both GFP and SiR channels was obtained, representing SiR-labelled and membrane-anchored 150BCNK-EGFP-CAAX (Fig. 4B). Under these conditions, SiR fluorescence intensity levels were significantly higher in the membrane than in the cytosol.

**Figure 4.**
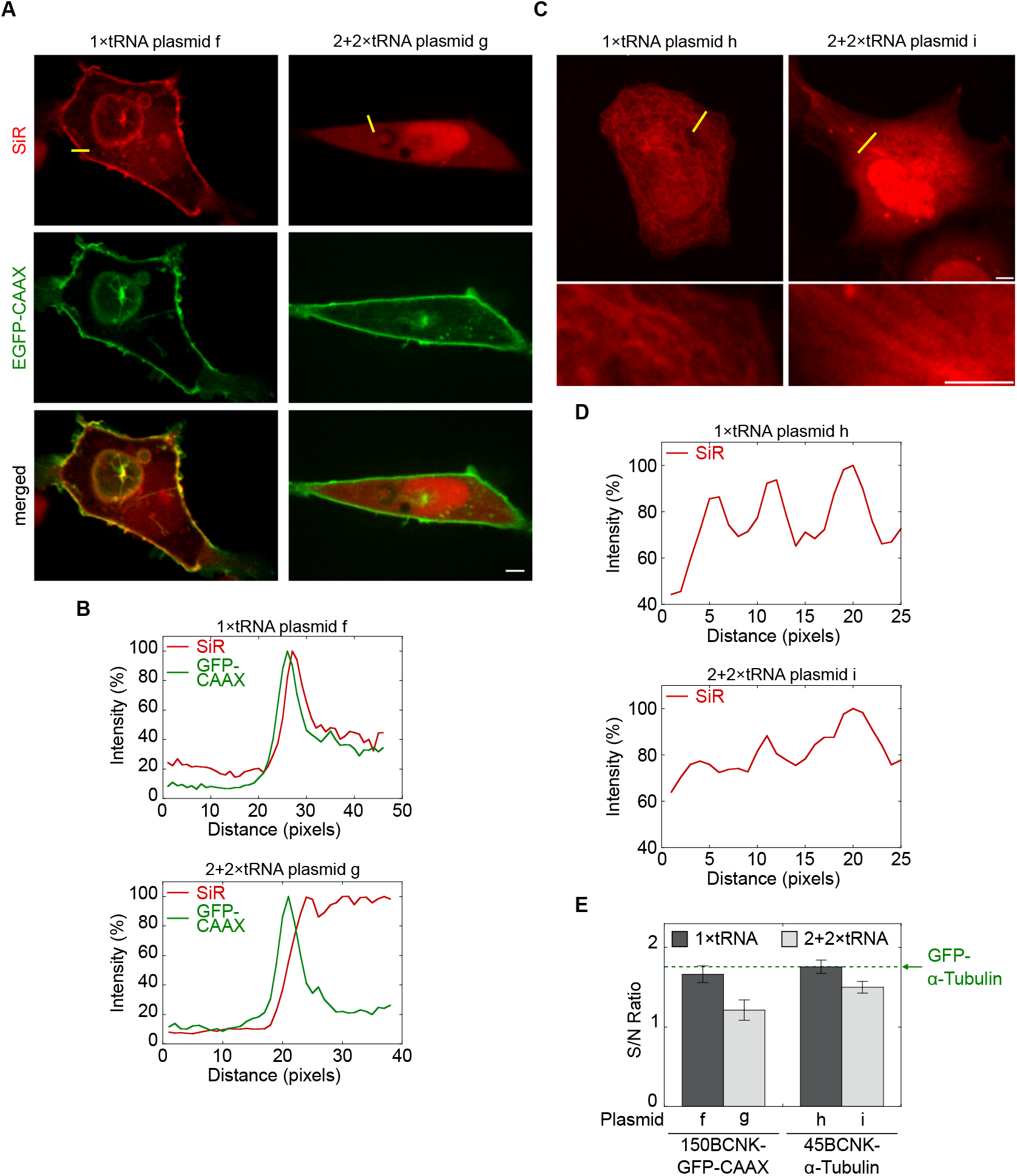
(**A**) Live cell imaging of SiR-labelled 150BCNK-EGFP-CAAX expressed in COS7 cells. Top row: red SiR; centre row: green EGFP; bottom row: merge. The scale bar represents 5 µm. (**B**) Line intensity profiles of 488 and 640 nm channels (EGFP and SiR, respectively) are presented as a percentage of the maximum value and are plotted as a function of distance along a line (yellow line in top panel A). (**C**) Live-cell imaging of SiR-labelled 45BCNK-α-tubulin expressed in COS7 cells. Zoomed-in images of a subset of the cell are presented below. The scale bar represents 5 µm. (**D**) Line intensity profiles of 640 nm channel presented as a percentage of the maximum value and plotted as a function of distance along a line (yellow line in top panel C). (**E**) Signal-to-noise ratios calculated from live-cell images of SiR-labelled 150BCNK-EGFP-CAAX and 45BCNK-α-tubulin expressed in COS7 cells using plasmids carrying 1 or 4 copies of PylT (*±* SEM). The green dashed line marks the signal-to-noise ratio of GFP-labelled α-tubulin visualized in live COS7 cells under identical conditions.

The use of a single copy of the encoded PylT gene instead of 4 also improved image quality of SiR-labelled 45BCNK-α-tubulin expressed in COS7 cells (Fig. 4C). In cells transfected with 2+2*×*PylT plasmid **i**, few and relatively thick SiR-labelled fibres could be seen in only a small subset of the transfected cells. In contrast, SiR-labelled microtubules were clearly visible in cells transfected with 1*×*PylT plasmid **h**. In addition, noticeable improvement in microtuble-labelling relative to cytosolic-labelling was observed in line intensity measurements (Fig. 4D). Overall, the improvement in image quality of SiR-labelled 150BCNK-EGFP-CAAX and 45BCNK-α-tubulin correlated with increases of 0.45 and 0.26 in signal-to-noise ratio, respectively (Fig. 4E). Relative to the signal-to-noise ratio of GFP-labelled α-tubulin visualized in live COS7 cells (1.75, Fig. 4E, green dashed line), the improvement in signal-to-noise ratio presented here is substantial.

### Reduction in pylT copy numbers correlates with a reduction in background fluorescence and in immobile tRNA^Pyl^ population

The SiR-labelling found in the cytosol, and more significantly in the nucleus (Fig. 4A and C), was ncAA-dependent (Supplementary Fig. S7) and was probably the result of a reaction between ncAA **2** and SiR-Tet. As mentioned above, such chemoselective and non-protein-specific background labelling is thought to be due to a reaction between tetrazine-conjugated fluorophores and aminoacylated tRNAs^40,42^. In light of the above described increase in signal-to-noise ratio (Fig. 4E), we characterized the effects of encoded PylT copy numbers on *overall* non-protein-specific labelling. For this, we created plasmids with 1 or 4 copies of the PylT genes and BCN-RS (plasmids **j** and **k**, respectively; Fig. 5A). Lacking a protein-of-interest-encoding gene with an in-frame TAG mutation, plasmids **j** and **k** allow for direct visualization and quantification of protein-independent background fluorescence resulting from SiR-Tet–labelled tRNA^Pyl^ without masking by a co-expressed labelled protein. Live-cell confocal microscopy using a highly sensitive single molecule setup, revealed that the averaged non-protein-specific fluorescence intensity in the nucleus and cytosol was reduced by *~*50% and *~*37%, respectively, when PylT copy numbers were reduced from 4 to 1 (Fig. 5B and C). Hence, reducing the number of encoded PylT genes improved the signal-to-noise ratio, by reducing background SiR-fluorescence. Furthermore, the similar decrease in fluorescence measured in the nucleus and cytosol confirmed that aminoacylated tRNA^Pyl^ was not entrapped in the nucleus, but rather was found in a state of dynamic equilibrium between the nucleus and cytosol.

**Figure 5.**
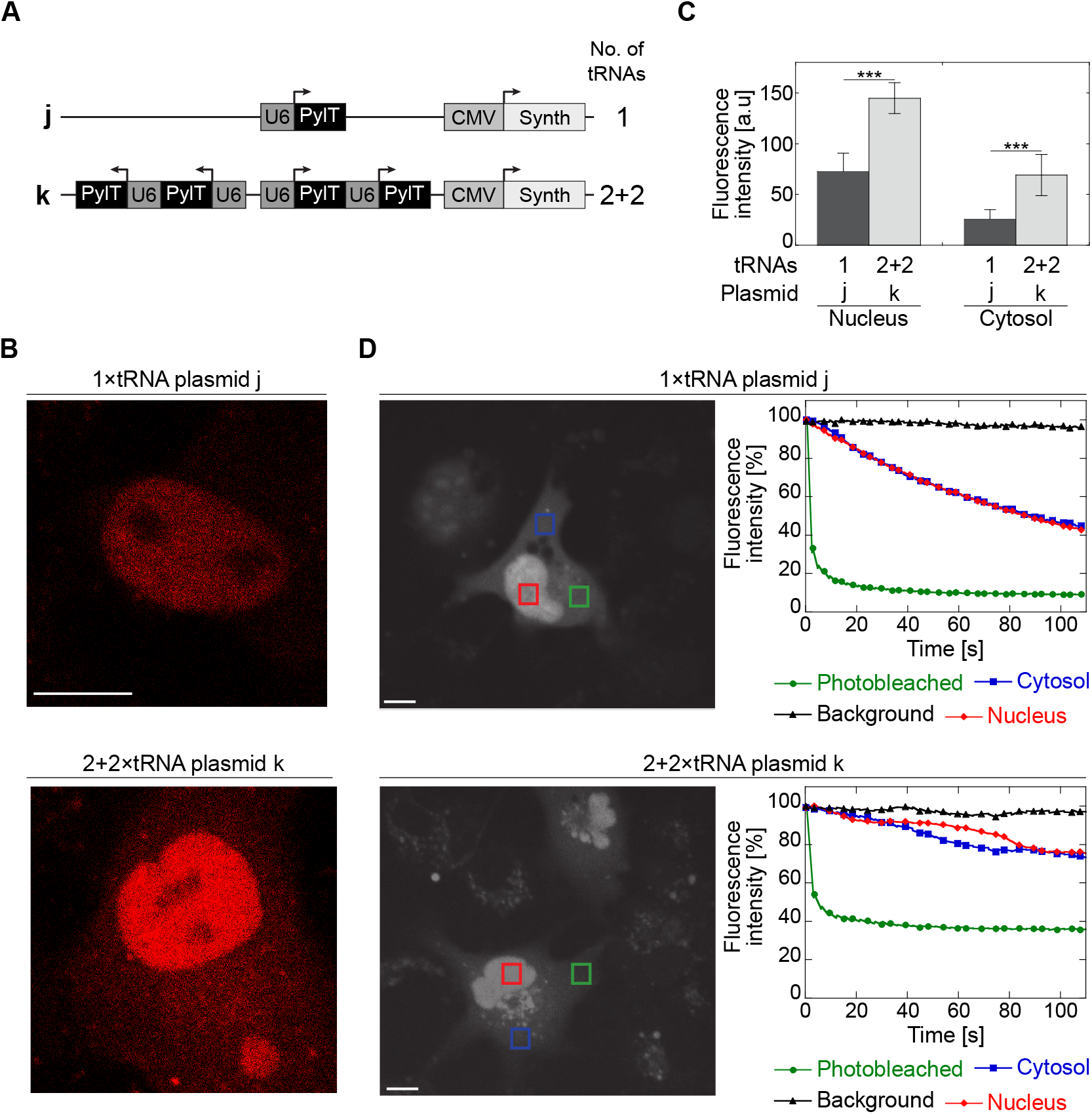
(**A**) Schematic representation of plasmids with 1 or 2+2 copies of PylT genes and BCN-RS used in the characterization of non-protein-specific labelling and measurement of aminoacylated tRNA^Pyl^ dynamics. (**B**) Non protein-specific background SiR fluorescence. COS7 cells transformed with plasmid **j** or **k**, labelled with SiR-Tet, and visualized by confocal microscope equipped with a single molecule detector. (**C**) SiR fluorescence measured in the nucleus (left) or cytosol (right) of COS7 cells treated as described in panel B. Values are displayed as average fluorescence intensity *±*SSD (n 7). Statistical analysis was performed using Student’s *t*-test, *** P *<*0.001. (**D**) FLIP measurements of COS7 cells described in panel B. Mean fluorescence intensity was measured in the photobleached area (green), the cytosol and nucleus of the photobleached cell (blue and red, respectively) as well as a nearby cell (background, black), and displayed as a function of time. The scale bar represents 10 µm.

We also noticed the presence of dense ‘fluorescent dots’ (Supplementary Fig. S8, white arrows) in many SiR-labelled cells transfected with a 2+2*×*PylT plasmid **k**, but not in cells transfected with 1*×*PylT plasmid j. We hypothesized that the distribution of tRNA^Pyl^ in the cell was non-homogeneous, especially when the tRNA was transcribed at high levels. To test our hypothesis, we measured the mobility of tRNA^Pyl^ in COS7 cells using fluorescence loss in photobleaching (FLIP), with the aim of identifying sub-populations of tRNA^Pyl^ entrapped in large complexes. In these studies, we compared SiR-Tet-labelled COS7 cells transfected with 1*×*PylT plasmid **j** and cells transfected with 2+2*×*PylT plasmid **k** (Fig. 5D). Over the course of two minutes, for a given number of encoded PylT genes, the time-dependent photobleaching pattern in the nucleus was similar to that seen in the cytosol, suggesting that the nuclear fraction of aminoacylated tRNA^Pyl^ was in equilibrium with the cytosolic fraction. This result was consistent with the similar reduction in nuclear and cytosolic background fluorescence observed upon decreasing PylT copy numbers (Fig. 5C). However, overall photobleaching efficiency was dependent on the number of encoded PylT genes. In cells transfected with 1*×*PylT plasmid **j** the fluorescence in the photobleached area was reduced to less than 10% and fluorescence in the cytosol and nucleus was reduced to 40% after two minutes (Fig. 5C, left panel). In contrast, when the same FLIP protocol was applied to cells transfected with 2+2*×*PylT plasmid **k**, fluorescence in the bleached area was reduced to only 40% and fluorescence in the cytosol and nucleus was only reduced to about 70% (Fig. 5C, right panel). These results suggest that in COS7 cells transiently transfected with the 2+2*×*PylT plasmid, there was an immobile and photobleaching-resistant population of aminoacylated and SiR-labelled tRNA^Pyl^. While this photobleaching-resistant population may have increased non-protein-specific background labelling, its origin and character remain to be determined.

### The use of one copy of pylT gene is sufficient for advanced imaging applications

In light of the observed improvement in signal-to-noise ratio following the reduction in the number of encoded PylT genes, we decided to use our single plasmid-based system to co-label two different proteins for live-cell fluorescence imaging. COS7 cells were co-transfected with 1*×*PylT plasmids **f** *and* **h**, or 2+2*×*PylT plasmids **g** *and* **i** (Fig. 3A) for co-expression and labelling of 150BCNK-EGFP-CAAX *and* 45BCNK-α-tubulin. Co-labelling experiments using 2+2*×*PylT plasmids **g** and **i** were, however, unsuccessful, as image quality was bellow the level obtained with single protein labelling. In contrast, live confocal microscopy showed clear SiR-labelling of both membrane-anchored 150BCNK-EGFP-CAAX and 45BCNK-α-tubulin in COS7 cells co-transfected with 1*×*PylT plasmids **f** and **h** (Fig. 6A). In addition, there was clear overlap between GFP fluorescence and SiR-labelling of the membrane. To the best of our knowledge, this is the first example of live-cell imaging of two different intracellular proteins bioorthogonally labelled with organic fluorophores obtained *via* genetic encoding of ncAAs.

**Figure 6.**
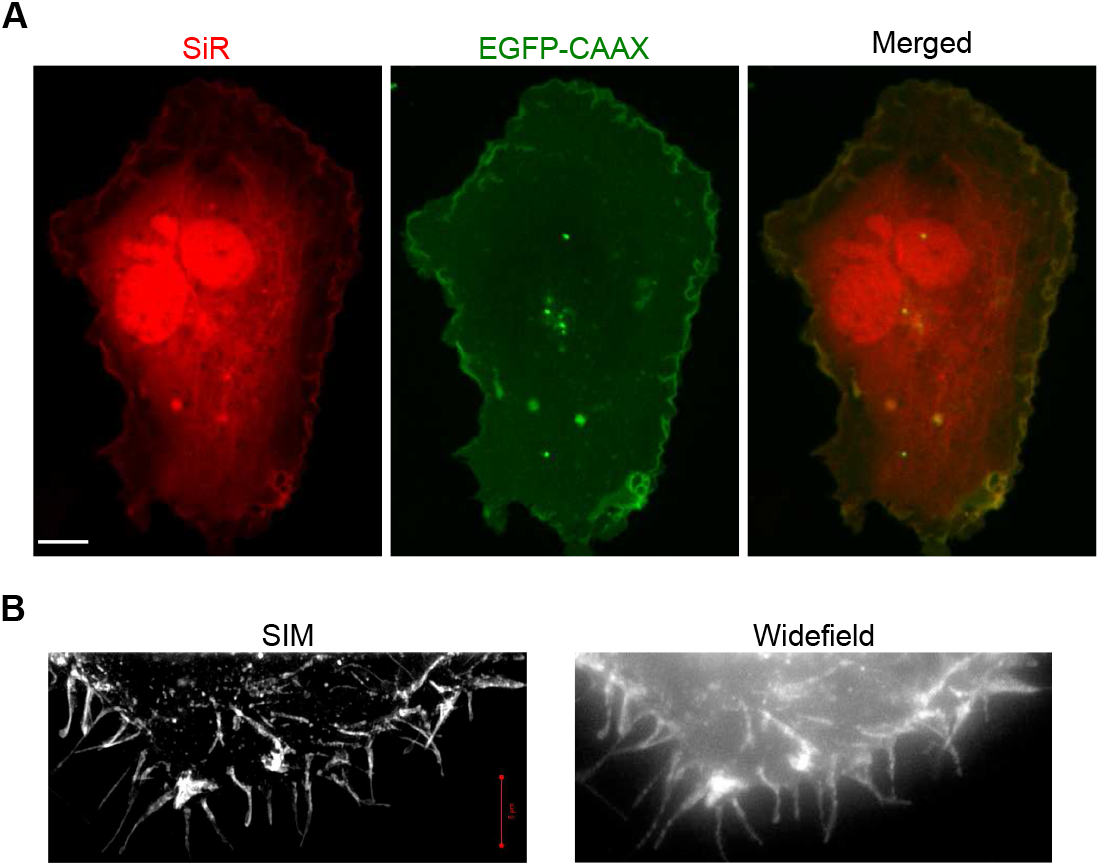
(**A**) COS7 cells co-transfected with plasmids **f** and **h**, incubated in the presence of **2**, and labelled with SiR-Tet. The use of a single copy of the encoded PylT gene enabled co-expression and SiR labelling of membrane-anchored 150BCNK-EGFP-CAAX and 45BCNK-α-tubulin. The scale bar represents 10 µm. (**B**) Zoomed-in regions taken from SIM and widefield imaging of SiR-labelled 150BCNK-EGFP-CAAX (Supplementary Fig. S9).

Finally, we visualized SiR-labelled 150BCNK-EGFP-CAAX using high-resolution structured illumination microscopy (SIM; Fig. 6B). COS7 cells were transfected with 1*×*PylT plasmid **f**, incubated with ncAA **2**, labelled with SiR-Tet and fixed before being visualized by SIM. Under these conditions, we were able to generate a 3D high-resolution image of the labelled membrane in SIM (Fig. 6B and Supplementary Fig. S9). This demonstrated that even with high resolution fluorescence imaging applications, such as SIM, one copy of encoded PylT gene is sufficient for efficient protein expression and subsequent bioorthogonal labelling of intracellular proteins.

## Discussion

Genetic code expansion is a powerful technology that enables the expression of tailor-made proteins by living organisms. When combined with site-specific and bioorthogonal protein labelling of proteins, genetic code expansion can have a strong impact on how we visualize proteins in living cells. That said, live-cell imaging applications have specific requirements that must be addressed. One important challenge encountered in the genetic encoding of a ncAA in response to an in-frame stop codon is low suppression efficiency and consequently, low protein expression levels. This challenge was addressed by several groups who were able to significantly improve our ability to efficiently incorporate ncAAs into proteins expressed in cultured mammalian cells. These studies also contributed to improving the fluorescence imaging of bioorthogonally labelled proteins. However, when it comes to addressing biologically relevant questions by visualizing intracellular proteins in living cells, maximal protein expression level is not the sole important parameter. In such applications, signal-to-noise ratio, proper processing and localization of the labelled protein, and minimal interference with cell physiology are equally important considerations that must be taken into account. With this in mind, we studied the effect of PylT copy numbers on image quality and signal-to-noise ratio, with the aim of improving these parameters, rather than protein expression yield, while minimizing interference with cell physiology.

It is well accepted that amber suppression efficiency correlates with intracellular tRNA^Pyl^ levels. We found that in COS7 cells transiently transfected with a single plasmid, increasing PylT copy numbers had only a negligible effect on amber suppression efficiency and overall protein expression levels. While the correlation between PylT copy numbers and cellular tRNA^Pyl^ levels has been previously reported, it is difficult to directly compare these studies due to differences in cell lines, the number of expression plasmids and promoter types used, DNA delivery methods, etc. (Supplementary Table S1). That said, Coin et al. recently found, in agreement with the data presented here, that the use of one U25C PylT gene rather than four had only a marginal and statistically insignificant effect on tRNA^Pyl^ levels in HEK293 cells^29^. Importantly, they found that tRNA^Pyl^ levels could be increased by using an engineered, and presumably more stable, PylT mutant. These results could explain our detection of a higher immobile tRNA^Pyl^ population in COS7 cells transfected with plasmids carrying four copies of PylT, suggesting that an excess of tRNA^Pyl^ was sequestered and had become unavailable. The identity of this immobile fraction, as well as those factors that promote high levels of available and functional tRNA^Pyl^ in mammalian cells, will be addressed in future studies.

It was recently argued that the fraction of nuclear tRNA^Pyl^ can be minimized through the addition of a nuclear export signal to PylRS and its evolved mutants^42^. However, any increase in the cytosolic tRNA^Pyl^ fraction may interfere with live-cell imaging of intracellular proteins using cell-permeable dyes due the an increase in cytosolic background fluorescence resulting from labelled tRNA^Pyl^. This is not a concern in imaging of fixed cells or in live cell imaging of extracellular proteins. It is, therefore, important to study how a shift in tRNA^Pyl^ equilibrium between the nucleus and cytosol affects signal-to-noise ratio and cytosolic background fluorescence in live-cell imaging applications using cell-permeable dyes.

In the present report, we improved image quality and signal-to-noise ratio in live-cell imaging of SiR-labelled proteins expressed in transiently transfected mammalian cells by reducing PylT copy numbers. The reduction in the number of encoded PylT genes resulted in reduced background fluorescence and was accompanied by an increase in the total number of cells with protein-specific SiR-labelling, following transfection and a multi-step labelling protocol. Hence, we suggest that for live-cell imaging of bioorthogonally labelled proteins using genetic code expansion technology and cell-permeable dyes, the number of encoded PylT genes should (and can be) minimized based on the cell line, the ncAA incorporated, and labelled protein.

## Acknowledgements

This work was funded by the European Research Council (ERC) under the European Union’s Horizon 2020 research and innovation programme under grant agreement No. 678461 (to E.A.) and No. 639313 (to N.E.), by the ARO under grant agreement No. 65422-LS (to N.E. and E.A.), and by the Israel Science Foundation (grant number 807/15 to E.A.).

## Author contributions statement

N.A., T.S., A.I.K., S.C., E.B., B.T., and D.N., conducted the experiments. N.A., D.N., Y.G., N.E., and E.A. analysed the results. O.B. provided technical assistance. N.A., and E.A. wrote the manuscript. E.A. conceived the experiments. All authors reviewed the manuscript.

